# Passage of time at the level of milliseconds : a new approach and a selective difficulty in individuals with schizophrenia

**DOI:** 10.64898/2026.01.09.698413

**Authors:** Ljubica Jovanovic, Laurence Lalanne, Florent Bernardin, Vincent Laprévote, Anne Giersch

## Abstract

**Background and Hypothesis:** Individuals with schizophrenia report that to them time sometimes feels discontinuous. Previous work has shown a link between the sense of self, and the ability to prepare to react to a target and to benefit from the passage of time, at the level of a few hundreds of milliseconds. However, the sense of time continuity requires a higher time resolution than examined in previous studies. Here we investigate to which extent individuals with schizophrenia and controls benefit from an increased delay when detecting asynchronies in the order of tens of milliseconds (stimulus onset asynchronies, i.e. SOA) between two successive visual stimuli.

**Study Design:** We re-analyzed three datasets and contrasted performance when the SOA increases, remains identical or decreases from trial t-1 to trial t.

**Study Results:** Performance of all participants improved with an increase in the SOA, as the task became easier, but for controls more so than individuals with schizophrenia. These results are replicated across datasets, and were specific to the condition when SOA increased from one trial to the next. There was no significant group difference when the SOA decreased or remained identical. This was true even when the latter condition was more frequent, and when SOA magnitude was equalized across the conditions. Importantly, the difference between the two groups was specific to temporal judgements, and was not observed in a control masking task.

**Conclusions:** We suggest the results reveal a difficulty for individuals with schizophrenia to experience time at the level of milliseconds.

---

Individuals with schizophrenia report that their conscious experience is fragmented in time, as if for them time itself would be discontinuous rather than continuous^1–3^. We explore the mechanisms associated with this disruption of time continuity, as it is believed to contribute to the disorders of the sense of self^1,4–6^. Consistent with the phenomenological descriptions, the passage of time is reported to be experienced as disturbed in individuals with schizophrenia. In former studies, the passage of time has been explored at the level of seconds or minutes^3,7^. However, disturbances have been also described at the level of milliseconds in individuals with schizophrenia^8–16^. We suggested that disturbances of the passage of time might be based on impairments in monitoring time at shorter time scales, specifically at the level of milliseconds^17,18^. If this were true, then alterations of the passage of time should be observed for shorter durations than one second in individuals with schizophrenia. In the present study we explore how the passage of time at the level of tens of milliseconds is used to temporally distinguish two successive visual events, in individuals with schizophrenia and in healthy controls.

In former studies, we tested the hypothesis of time disruption experimentally, by evaluating the capacity to exploit the passage of time when anticipating events. To that aim we used the variable foreperiod paradigm. Participants responded to a target presentation by pressing on a response key. The target occurred at variable delays (foreperiods) after a warning signal. Typically, control participants get better prepared as time passes and react faster to the target when foreperiods are long rather than short^19^. We showed that individuals with schizophrenia did not benefit from the passage of time as much as the control participants, and that these disturbances are correlated with disorders of the sense of self^20,21^.

However, it is difficult to explore our hypothesis of an impaired time monitoring at the millisecond time scale with the variable foreperiod paradigm, as it is difficult to distinguish and react to distinct successive stimuli in time when their stimulus onset asynchrony (SOA) is less than 20-50 ms ^22^. To address the question of the implicit passage of time at the subsecond scale, we developped a different approach. We used a task requiring participants to detect and report an asynchrony between the presentation of two successive visual stimuli. The presented SOAs varied from trial to trial, and we investigated sequential effects : how the SOA and response on the previous trial t-1 affect performance on the current trial t. Specifically, we tested the possibility that participants take advantage from the increase in SOA from one trial to another (e.g. for trial t-1 SOA = 10 ms and for trial t SOA=40 ms) to improve their performance (Fig 1). Of course, an increase in SOA from one trial to the next makes the task easier and an improvement in the performance is expected. However, there may be more to this improvement, if sensory signals are monitored over time at the level of milliseconds^23^. If the passage of time is monitored implicitly at the millisecond level and continues automatically beyond the delay of the previous trial, then the SOA increase from one trial to the next may be monitored as such, and would further facilitate the detection of an asynchrony. If the passage of time is not monitored efficiently at the millisecond level, only the benefit of task ease would remain.

**Figure 1.**
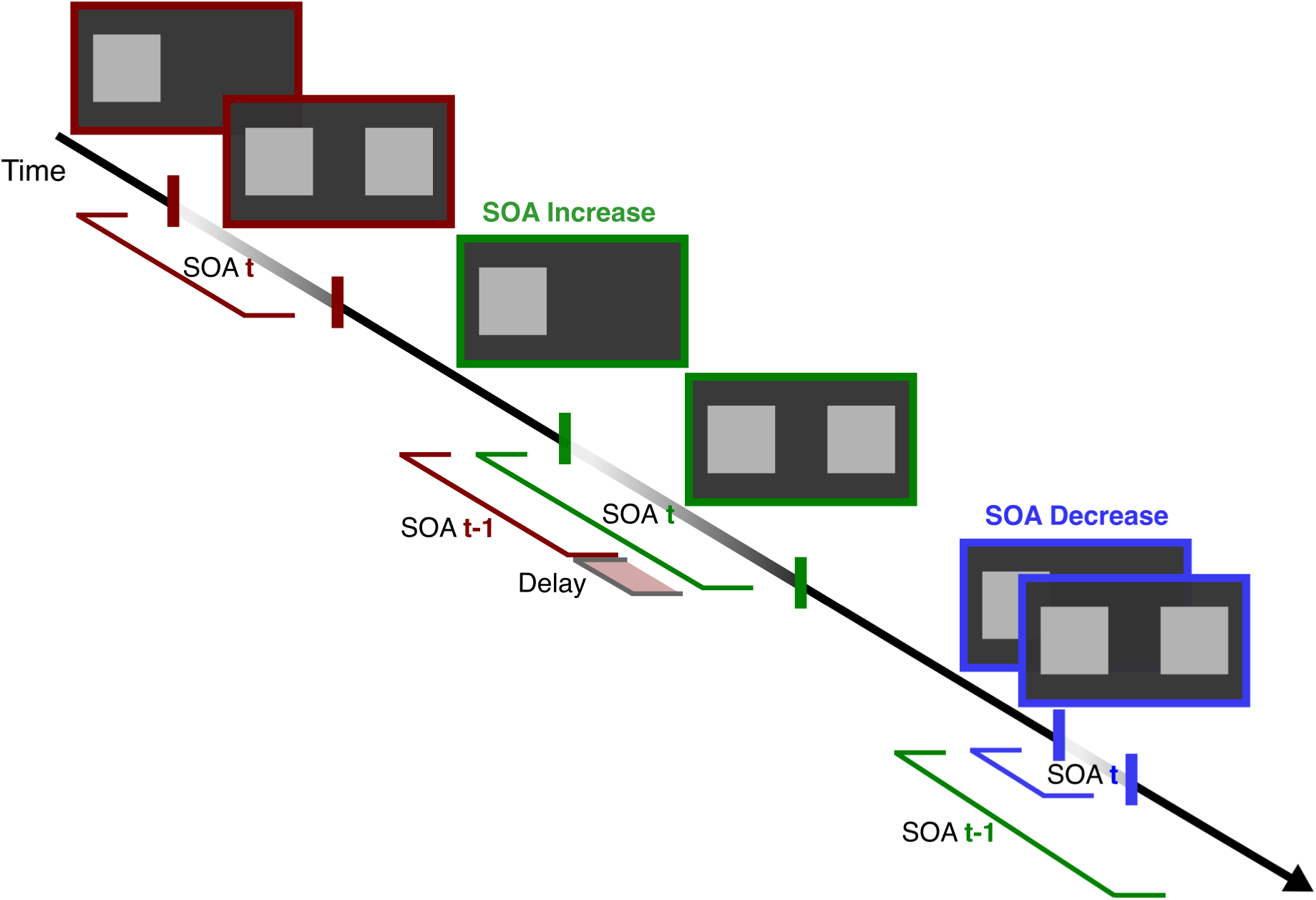
Illustration of the changes in SOA from trial to trial for the increasing (from red to green) and decreasing (from green to blue) conditions.

Given the difficulty of the individuals with schizophrenia to benefit of the passage of time^20,21^, and our hypothesis that their difficulty originates in impairments at the level of milliseconds, in this group we expect only a benefit of the increase in task easiness. In contrast, controls may display greater improvement when SOA increases. Moreover, a benefit of the passage of time should be observed specifically in case of a SOA increase (Fig 1). When the SOA remains the same, SOA expectation can be based on the previous trial. Similarly, when the SOA decreases there can hardly be a benefit of the passage of time. If individuals with schizophrenia have aspecific impairment in monitoring the passage of time at the level of milliseconds, an impairment relative to controls should be observed only in the case of a SOA increase.

## Methods

### Participants

We re-analyzed datasests from 2 published studies testing visual asynchrony discrimination (Study 1 and 3 in the manuscript), and one yet unpublished dataset (Study 2). In Study 1, 20 controls and 20 individuals with schizophrenia participated^13^. In Study 2, 29 controls and 26 individuals with schizophrenia were included. In Study 3, 18 participants were tested per group^12^, In addition, to test the specificity of the observed effects, we re-analyzed data from a masking experiment ^24^ (Study 4), in which 19 controls and 19 individuals with schizophrenia were tested. Demographic and clinical characteristics are summarized for each sample in Table 1.

**Table 1.**
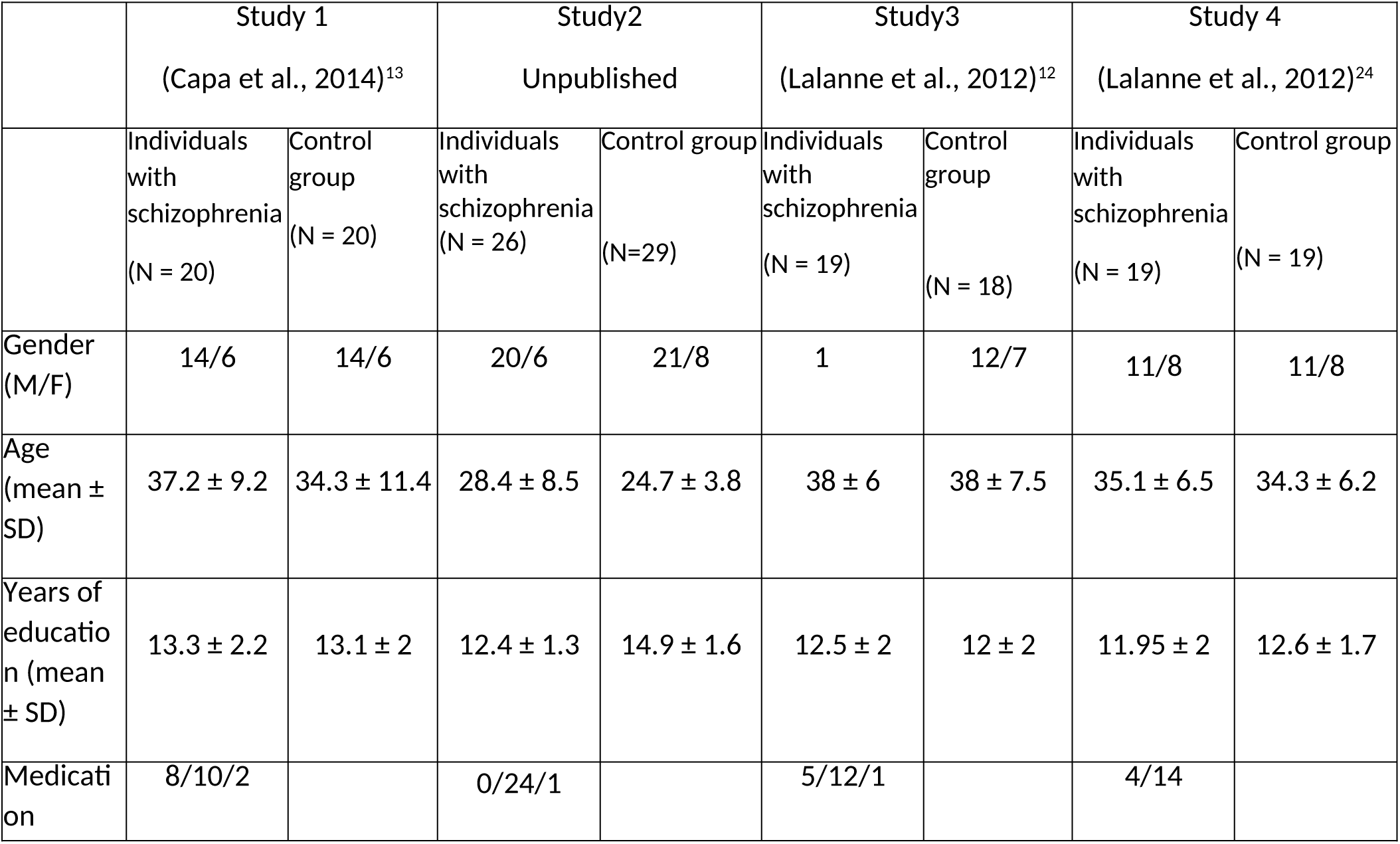
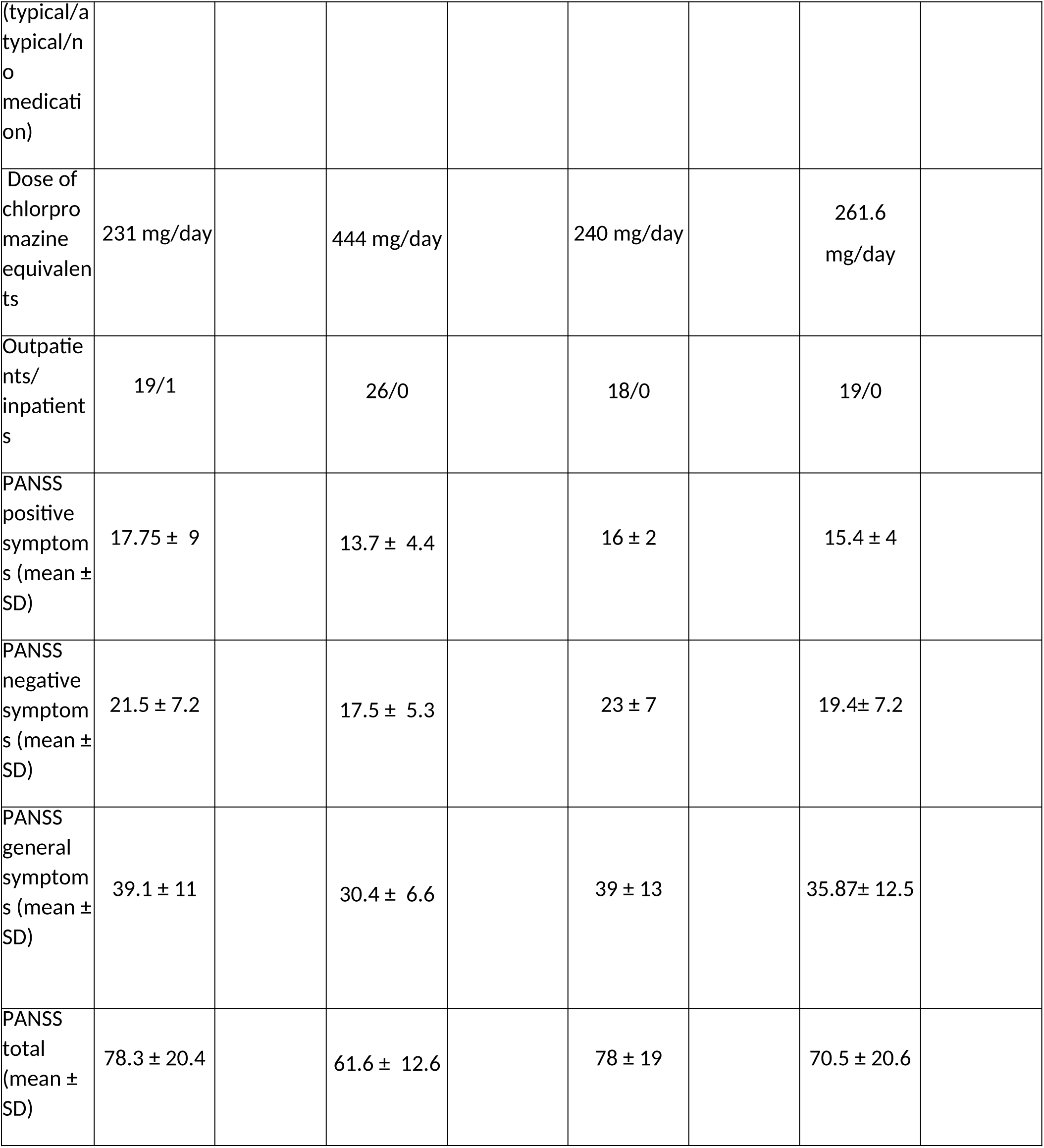
Demographic and clinical data of the participants in the four experiments.

|  | Study 1<br>(Capa et al., 2014)<sup>13</sup> |  | Study2<br>Unpublished |  | Study3<br>(Lalanne et al., 2012)<sup>12</sup> |  | Study 4<br>(Lalanne et al., 2012)<sup>24</sup> |  |
| --- | --- | --- | --- | --- | --- | --- | --- | --- |
|  | Individuals with schizophrenia<br>(N = 20) | Control group<br>(N = 20) | Individuals with schizophrenia<br>(N = 26) | Control group<br>(N=29) | Individuals with schizophrenia<br>(N = 19) | Control group<br>(N = 18) | Individuals with schizophrenia<br>(N = 19) | Control group<br>(N = 19) |
| Gender (M/F) | 14/6 | 14/6 | 20/6 | 21/8 | 1 | 12/7 | 11/8 | 11/8 |
| Age (mean ± SD) | 37.2 ± 9.2 | 34.3 ± 11.4 | 28.4 ± 8.5 | 24.7 ± 3.8 | 38 ± 6 | 38 ± 7.5 | 35.1 ± 6.5 | 34.3 ± 6.2 |
| Years of education (mean ± SD) | 13.3 ± 2.2 | 13.1 ± 2 | 12.4 ± 1.3 | 14.9 ± 1.6 | 12.5 ± 2 | 12 ± 2 | 11.95 ± 2 | 12.6 ± 1.7 |
| Medication | 8/10/2 |  | 0/24/1 |  | 5/12/1 |  | 4/14 |  |
| (typical/a<br>typical/n<br>o<br>medicati<br>on) |  |  |  |  |  |  |  |  |
| Dose of<br>chlorpro<br>mazine<br>equivalen<br>ts | 231 mg/day |  | 444 mg/day |  | 240 mg/day |  | 261.6<br>mg/day |  |
| Outpatie<br>nts/<br>inpatient<br>s | 19/1 |  | 26/0 |  | 18/0 |  | 19/0 |  |
| PANSS<br>positive<br>symptom<br>s (mean ±<br>SD) | 17.75 ± 9 | | 13.7 ± 4.4 | | 16 ± 2 | | 15.4 ± 4 | |
| PANSS<br>negative<br>symptom<br>s (mean ±<br>SD) | 21.5 ± 7.2 | | 17.5 ± 5.3 | | 23 ± 7 | | 19.4 ± 7.2 | |
| PANSS<br>general<br>symptom<br>s (mean ±<br>SD) | 39.1 ± 11 | | 30.4 ± 6.6 | | 39 ± 13 | | 35.87 ± 12.5 | |
| PANSS<br>total<br>(mean ±<br>SD) | 78.3 ± 20.4 | | 61.6 ± 12.6 | | 78 ± 19 | | 70.5 ± 20.6 | |

### Apparatus and stimuli

Apparatus and stimuli differed slightly across the studies, and are shown in Table 2.

**Table 2.** Stimuli and apparatus.

|  | Study 1 | Study 2 | Study 3 | Study 4 |
| --- | --- | --- | --- | --- |
| Monitor (refresh rate) | liyama 21 inches (85 Hz) | EIZO 21 inches (120 Hz) |  |  |
| Stimuli | 3.6 x 1.1 rectangles presented at 9.8° eccentricity, 12 cd/m<sup>2</sup> on background luminance 0.02 cd/m<sup>2</sup> | 0.8 x 0.8 squares at 5.5 deg of horizontal and vertical eccentricity. 12 cd/m<sup>2</sup> on background luminance 0.02 cd/m<sup>2</sup> | 0.8 x 0.8 squares at 5.5 deg of horizontal and vertical eccentricity. 12 cd/m<sup>2</sup> on background luminance 0.02 cd/m<sup>2</sup> | 0.8 x 0.8 squares at 5.5 deg of horizontal and vertical eccentricity. 3.26 cd/m<sup>2</sup> on background luminance 0.02 cd/m<sup>2</sup><br>The mask was presented for 33 ms with same luminance as the targets. |
| SOA (ms) | 0, 24, 48, 72, and 96 | 17,33,50,67,83 ms | 0, 8, 17, 25, 33, 41, 50, 58, 67, 75, 83, 92 ms | -250 to +250 ms by step of 50 ms |
| Number of trials per condition | 108 trials /SOA | 48 trials / SOA<br>48 trials in which the SOA remained identical with the same direction (left-right or right-left) in trials t-1 and t. | 64 trials / SOA | 24 trials / SOA |
|  | Pentium 4 PC |  |  |  |

Stimuli presentation was controlled by a custom-build software using Psychtoolbox^25^ (Study 1) and Cambridge Research System (Rochester, Kend, UK; Study 2, 3 and 4) running with Matlab 7.0.1 (Mathworks, 1984-2004).

### Procedures

The simultaneity/asynchrony discrimination task (Study 1-3) was the same in the three studies. On each trial, two stimuli were displayed either simultaneously or with an asynchrony below 100 ms, and participants decided whether or not the stimuli were asynchronous. They gave their response by pressing the ‘f’ key in case of a simultaneity, and the ‘j’ key in case of an asynchrony. The location of the stimuli and the precise SOAs differed between experiments, and are detailed in Table 2.

In Study 1, all participants first completed a temporal order judgement task (not shown here), before completing the asynchrony discrimination task that is presented in this manuscript. In Study 2, participants completed a typical asynchrony discrimination task, but there was no simultaenous condition. Furthermore, the SOA and stimuli direction (left-right or right-left) remained the same in the two consecutive trials in 48 trials. This allowed us to verify whether these repetitions improved the performance more so than a SOA increase from one trial to the next. In both Studies 1 and 2, there were 2 lateralized stimuli, one to the left and one to the right from the central fixation point. In Study 3 and 4, four empty squares were drawn in the four corners of a virtual square (5.5°x 5.5°) around a central fixation point. In a quarter of trials connectors linked squares horizontally, and in a quarter of trials, they linked squares vertically. In the remaining trials there was no connector. On each trial only two squares were filled in, at the top, bottom, or left or right side of the virtual square. In the asynchrony discrimination task in Study 3, the two squares were displayed with various SOAs like in Studies 1 and 2 (Table 2). Study 4^24^ was designed to test forward and backward masking on the detection of visual stimuli. On each trial, two targets were presented in the same locations as in Study 3, but always simultaneously. The targets were preceeded (forward masking) or followed (backward masking) by a mask, consisting of four squares presented at each of the four tested conditions. In Study 4, participants were asked to locate the target.

### Analyses

To test the hypothesis of the implicit monitoring of the passage of time at the millisecond level, we tested the effects of previous trials (t-1) on performance on the current trial (t), for the two groups of participants. More precisely, we tested whether the performance improved when there was an increase in SOA from one trial to the next one, as a function of their response on the previous trial (correct or incorrect). Responses on trials t-1 were considered separately, assuming that incorrect responses reflected an incorrect detection of the asynchrony. Furthermore, we considered trials with simultaneous squares as a special case, as simultaneity may elicit special processing, e.g. by the detection of coincidence^26–28^. We thus reasoned that the transition between a trial with simultaneous squares to asynchronous squares may differ from a transition from one given asynchrony to another. For this reason we discarded all trials with simultaneous squares (on the current or on the previous trial). In all the analyses we compared the SOA on trial t-1 and on trial t. We then grouped performance across trials as a function of the response on trial t-1 (asynchrony correctly detected, or not), and of the SOA change from trial t-1 to trial t (decrease, no change, increase in SOA). Note that this procedure grouped SOAs of different magnitudes, as trials preceded by a trial with smaller SOAs necessarily include larger SOAs than trials preceded by a trial with larger SOAs. Given the manner in which SOA direction is defined, the three groups were different with respect to SOA on trials t for the three groups. Specifically, in the “increasing” group there were no trials with the smallest, and in the “decreasing” group no trials with the largest SOA (see Fig. S1). Here we present the analysis on the data where the two extreme SOAs for each experiment were excluded, in order to match the difficulty across the categories. We have also repeated the analysis without the exclusion of the extreme SOAs for each experiment (Fig. S2). The detailed results of this analysis are shown in the Supplementary Information. Finally, we also repeated the analysis with inclusion of SOA on the current trial (t), shown in the Supplementary Information. Importantly, in both analyses the key findings were replicated.

## Results

### Studies 1-3: Sequential effects in asynchrony judgments

Performance on trial t is shown in Figure 1 as as a function of the three categories of SOA difference between t-1 and t, separately for correct and incorrect decisions on the previous trial (columns). Each row shows performance in a different study (Study 1-3), and color codes the two groups (control group and individuals with schizophrenia). As shown in Figure 1, performance was better when SOA increased from trial t-1 to t, compared to conditions when it was the same or decreased. This was true regardless of the response on the previous trial, and in the three studies. Of note, trials on which SOA increases are easier than previous trials, which in part explains the increase in the performance. However, if individuals with schizophrenia have difficulties in benefiting from the passage of time at these short timescales, we expect their improvement to be smaller than that of individuals in the control group.

**Figure 2.**
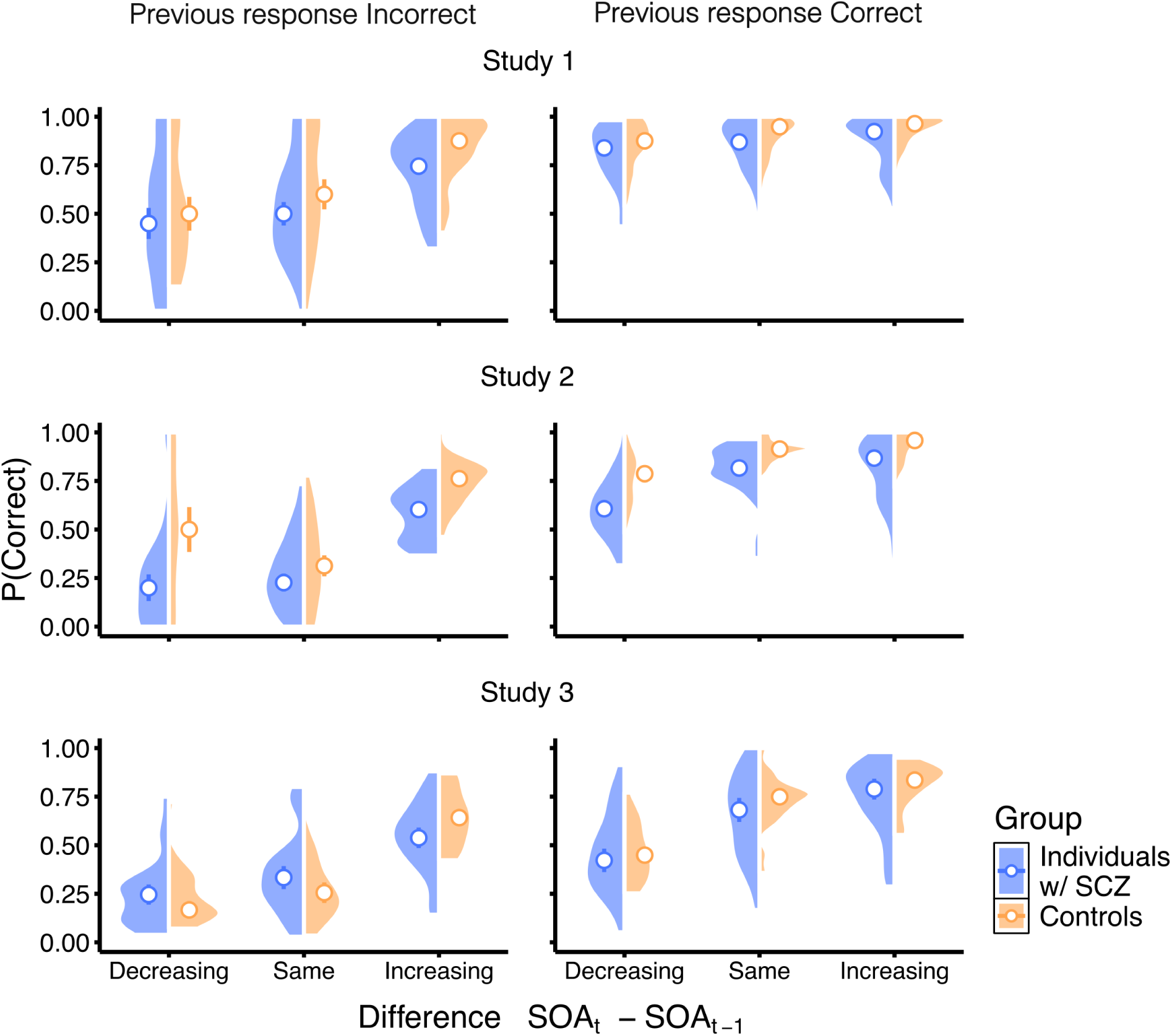
Sequential effects in the asynchrony detection task. Proportion of correct responses is plotted as a function of the difference between the SOA on the previous (t-1) and current (t) trial. The differences between the two consecutive trials are grouped in three groups, with respect to direction of SOA change (Decreasing, Same or Increasing). The left and right columns show performance when response on the previous trial (t-1) was incorrect and correct, respectively. The rows show performance for the three analysed datasets. Color codes the group, with blue symbols indicating performance of individuals with schizophrenia and orange that of individuals in the control group. Violin plots show distributions of the proportions of correct responses, for the two groups. Circles show median performance (error bars indicate standard error of the median across the participants).

In order to test our hypothesis simultaneously with all three datasets, we used a generalised linear-mixed effect model. Log-odds of correct response were predicted by the fixed factor structure consisting of the group (controls or individuals with schizophrenia), the study (1-3), the difference in SOA between the two successive trials (decreasing, same or increasing), response on the previous trial (correct or incorrect), and their interactions. A random intercept was specified at the level of participant, nested within the study. However, given the complexity of the model, the full model ran into convergence issues. In order to automate identification of the maximal generalised linear model that converges given the data, we used buildmer function (see Supplementary Information for details^29,30^). Odds ratios (OR) are shown as measures of effect size^31^.

In particular, there was a significant effect of the SOA direction (χ^2^(2) = 555.321, p<0.01), the response on the previous trial (χ^2^(1) = 304.433, p<0.01), and the study (χ^2^(2) = 48.289, p<0.01). Relative to the condition where the SOA decreased from trial t-1 to trial t, the performance was better when the SOA increased (β_SOA Increasing_ = 1.205, SE = 0.058, z = 20.635, p < 0.01, OR = 3.336; β_SOA Same_ = 0.170, SE = 0.058, z = 2.199, p < 0.05, OR = 1.19). Furthermore, when the response on the previous trial was correct, log-odds of correct response on trial t were greater (β_Response_t-1=Correct_ = 1.403, SE = 0.089, z = 17.448, p < 0.01, OR = 4.067). Finally, there was an effect of the study, and relative to the Study 1, overall performance was worse in Study 2 and Study 3 (β_Study2_ = −0.867, SE = 0.169, z = −5.111, p < 0.01, OR = 0.420; β_Study3_ = −1.120, SE = 0.182, z = −6.553, p < 0.01, OR = 0.326).

There was no evidence for a significant main effect of the group (χ^2^(1) = 2.2, p=0.137). Importantly, the group interacted with the SOA direction (χ^2^(2) = 60.484, p<0.01), such that performance of participants in the control group increased more when the SOA increased from the previous to the current trial, relative to individuals with schizophrenia (β_SOA Increasing x Controls_ = 0.487, SE = 0.066, z = 7.411, p < 0.01, OR = 1.627; β_SOA Same x Controls_ = 0.060, SE = 0.080, z = 0.727, p = 0.467, OR = 1.06).

The SOA direction also interacted with the previous response (χ^2^(2) = 108.887, p<0.01. The interaction with the previous response indicated that an increase in the performance for trials with a relative SOA increasing was smaller when response on the previous trial was correct (β_SOA Increasing x Response_t-1=Correct_ = −0.185, SE = 0.069, z = −2.671, p < 0.01, OR = 0.831). When relative SOA was the same, the effect was greater if the previous response was correct; β_SOA Same x Response_t-1=Correct_ = 0.676, SE = 0.086, z = 7.862, p < 0.01, OR = 1.966).

There was also an interaction between response on the previous trial and the study (χ^2^(2) = 128.2476, p<0.01), suggesting that the improvement in performance when response was correct on the previous trial was smaller in Studies 2 and 3 (β_Response_t-1=Correct x Study3_ = −0.570, SE = 0.07, z = −8.098, p < 0.01, OR = 0.565; β_Response_t-1=Correct x Study3_ = 0.09, SE = 0.082, z = 1.158, p = 247, OR = 1.09). Finally, there was evidence for an interaction between the response on the previous trial and the group (χ^2^(1) = 8.535, p<0.01), such that the effect when the previous response was correct was greater in the control group (β_Response_t-1=Correct x Controls_ = 0.181, SE = 0.07, z = 2.922, p < 0.01, OR = 1.198).

In support of our hypothesis, there was evidence for a greater improvement in performance for the increasing SOA condition in the control group of individuals compared to individuals with schizophrenia.

### Study 4: Masking experiment

Results presented so far indicate that individuals with schizophrenia do not improve their performance as much as the participants in the control group, when the SOA is longer on trial the current than on the previous trial. These results are in line with the hypothesis that individuals with schizophrenia have difficulties to implicitly use the passage of time at the level of milliseconds. However, it is possible that they simply benefit less from the task becoming easier, regardless of the nature of the task. They may also have difficulties in segregating the successive stimuli, thus making it more difficult to benefit from the SOA increase. This segregation does not necessary entail a representation of the delay between the successive stimuli, as asynchrony and duration are two orthogonal concepts^17^. To test the possibility of a temporal segregation impairment, we analysed sequential effects in a masking paradigm, in which participants locate a target presented at various delays before or after a mask. Segregating the two successive stimuli helps in both the masking and the asynchrony detection tasks, but representing the delay between the two successive stimuli may further help the detection of an asynchrony : if a delay is detected between the two successive stimuli, it necessarily means an asynchrony. In contrast, representing the delay is of no use in the masking task : whether the delay is detectable as such, or not, does not help to locate the target. If the coding of the delay is impaired in individuals with schizophrenia, then a group effect should be observed selectively in the asynchrony detection task, whereas if it is the segregation of the stimuli in time that is impaired, their performance should be impaired in both tasks.

To test these alternatives, we re-analyzed a dataset from a study with a different task, with a design closely matching the visual display of Study 3 presented in this manuscript. Briefly, it was a novel masking paradigm, with two targets instead of only one typically presented in masking paradigms^24^. We chose this task because its spatial complexity required a high number of trials, thus allowing for a calculation of performance as a function of the SOA change between trials.

We considered consecutive trials without order change, in which both trials had been with forward or with backward masking, and consecutive trials with order change, in which trial t-1 had been with forward masking and trial t with backward masking, or the reverse. As a matter of fact the mechanisms underlying the two types of masking are suggested to differ^32^, which could have in turn precluded any benefit of SOA increase.

To test the specificity of the sequential effects obtained in the simultaneity/asynchrony discrimination task, we fitted data to a generalised linear mixed-effect model, with response on trial t as dependent variable, and the four fixed effects (group, response on trial t-1, the difference in SOA, the difference in type of masking between the two consecutive trials) and their interactions as predictors. Participants were included as random intercept. We used the same analyses as before, with automatised model selection. As before, we included only intermediate SOAs, shared between the conditions. For analysis on the full dataset, see Supplementary Information. The final model consisted of the fixed effects for the four predictors, two-way interactions between the previous response and the difference in SOA, type of masking and group and a three-way interaction between the response on trial t-1, the masking type and group. We found an effect of the response on the previous trial (χ^2^(1) = 9.927 p<0.01), and responses were more likely to be correct if response on the previous trial was correct as well (β_Previous Correct_ = 0.357, SE = 0.113, z = 3.145, p < 0.01, OR = 1.429). There was also an effect of the difference in SOA in the two consecutive trials (χ^2^(2) = 12,756 p<0.01), and performance was better when SOA on the current trial t was greater than that of the previous trial t-1 (β_SOA Incrasing_ = 0.229, SE = 0.064, z = 3.567, p < 0.01, oR = 1.257; β_SOA Same_ = 0.1101, SE = 0.083, z = 1.207, p = 0.227, OR = 1.116), but there was no evidence for an interaction with the group. There was also evidence for a three-way interaction between the group, the response on the previous trial and the relative masking type (χ^2^(1) = 10.604, p<0.01). These effects suggest that the effects of the previous response in the control group depended on the difference in the masking type from one trial to the other (β_Previous Correct x Control x Masking Same_ = 0.759, SE = 0.233, z = 3.254, p < 0.01, OR = 2.136). Similar results were obtained when we considered only the trials in which the masking type did not change from one trial to the next (see Supplementary Information for more details and visual representation of results). In summary, the pattern of results observed here is in agreement with the hypothesis that individuals with schizophrenia have reduced sensitivity to passage of time at the milliseconds level rather than a difficulty to benefit from the task becoming easier, or a non-specific difficulty to segregate stimuli in time.

### Correlations

We correlated the change in performance across conditions with clinical parameters, including the dosage of antipsychotics, but did not find any correlation.

## Discussion

Not surprisingly, the results show that all participants benefit from the task becoming easier from one trial to the next. This was observed in two types of tasks. First, it was observed when participants were asked to detect an asynchrony between two successive stimuli shown in distinct locations (Studies 1-3). Second, the benefit of the task becoming easier was observed in the masking experiment, when participants detected whether or not a target stimulus had been shown just before or just after a mask displayed in the same location as the target stimulus (Study 4). Most importantly, the improvement observed with the SOA increase was larger in the control group than in individuals with schizophrenia, and this improvement was task specific. It occurred only in the simultaneity/asynchrony discrimination task, not in the masking task. The results are consistent with our hypothesis that the individuals in the control group benefit from the passage of time at the level of milliseconds when detection of asynchronies is important for the task, beyond the advantage provided by the fact that the task was easier. Nevertheless, some alternatives need to be discussed, before setting on that conclusion.

The fact that in the masking task all participants benefit equally from the task becoming easier is important. This result was obtained in a similar way to the individual analyses of each asynchrony detection task set (analyses in Supplementary Information). This result indicates that we can exclude an explanation of the results on the temporal task in terms of a non specific insensitivity to the task becoming easier, and that the improvement is related to the nature of the task. Most importantly, the masking results also confirm that individuals with schizophrenia are sensitive to the SOA increase from trial to trial. This may appear as surprising, given their difficulties to explicitly detect asynchronies documented in the literature^8–16^. However, despite the difficulty of individuals with schizophrenia to explicitly report asynchronies, several studies indicate that they are implicitly affected by short asynchronies, and sometimes more so than controls^11,12^. Here we show that in spite of their sensitivity to short asynchronies, individuals with schizophrenia do not benefit from this increase as much as controls when the task is to detect an asynchrony. Is there another explanation for this group difference than a difficulty to benefit from the passage of time?

First it should be noted that the results obtained in the present study in the simultaneity/asynchrony tasks cannot be easily discarded. They are replicated in three datasets, with individuals with schizophrenia from different centres (in Strasbourg or in Nancy). The group difference was selectively observed when the SOA increased between trial t-1 and trial t. One alternative explanation is that this result is a consequence of the manner in which SOAs were grouped for the analysis. When SOAs increase, and the SOAs on trial t are around threshold, then SOAs on trial t-1 are sub-threshold, and individuals with schizophrenia have difficulty in processing sub-threshold asynchronies^11,12^. Several studies have shown these sub-threshold asynchronies to be processed even when not explicitly reported^11,12,33^. Participants in the control group tend to press to the side of the second stimulus of the sequence after the display of the two stimuli, even for small asynchronies. In contrast, individuals with schizophrenia tend to press to the side of the first stimulus at sub-threshold asynchronies, as if stuck with the first stimulus of the sequence^11,12^. Furthermore, it might have been suggested that individuals with schizophrenia do not predict or follow sub-threshold sequences on trial t-1, and this is why it is more difficult for them to benefit from the SOA increase on trial t (going from shorter to longer asynchronies from one trial to the next). However several results contradict this possibility. The same pattern of results, i.e. a lack of benefit of the SOA increase in individuals with schizophrenia, are observed when all SOAs are taken into account (i.e. when the shortest asynchronies are also included; see Supplementary Information). Moreover, had it been a difficulty for them to predict an asynchrony on the basis of the previous trial, then a group difference should have been observed also when the SOA stayed the same. However there was no group difference in this condition, whether with rarely or frequently repeated SOAs. Therefore, it seems unlikely that the lack of the benefit of the SOA increase in individuals with schizophrenia is due to a difficulty in predicting or following the current SOA on the basis of the previous trial. The results are rather consistent with the prior observations that individuals with schizophrenia are disturbed when the delay on trial t is not exactly identical to the delay in trial t-1, at least when those delays are small^34^.

As a matter of fact, when participants made pointing actions on a virtual surface, and the haptic feedback (i.e. the moment of the contact with the surface) was postponed, the feeling of control of individuals with schizophrenia decreased even when the delay was only 15 ms, whereas the feeling of control remained stable in the control group for short delays. In the context of this study, we might expect individuals with schizophrenia to be perturbed when the SOA increases because they are oversensitive to the SOA changes from trial to trial. This might explain a smaller benefit from the SOA increase in individuals with schizophrenia relative to controls. Two results do not fit with this hypothesis. First, a reduction of SOA should perturb their performance in the same manner, which was not the case. Additionally, if the individuals with schizophrenia were implicitly perturbed by a SOA increase, then we would expect the same pattern of results in the masking experiment (Study 4), which is not what we observed.

In fact, the contrast between the results found in the masking and the asynchrony detection task is consistent with our hypothesis of a decreased benefit of the passage of time in individuals with schizophrenia. As already emphasized, one important difference between the masking and simultaneity/asynchrony discrimination task concerns the implicit vs. explicit impact of the SOA. In the masking task, participants have to localize a target whose visibility is reduced by means of a mask displayed in close temporal succession. Time intervenes incidentally by increasing or decreasing the visibility of the target, but individuals do not have to make any explicit temporal judgement. In contrast, in the simultaneity/asynchrony discrimination task, participants have to judge whether stimuli are separated in time or not. Asynchrony detection is closely related to the existence of a time interval in-between the two stimuli onsets. Hence detecting the delay is a way to detect the asynchrony. In the present study, what makes the difference between the two tasks (masking vs. asynchrony detection) is the use of the additional delay in the explicit detection of asynchrony. In other words, the results suggest that the increase in the delay from trial t-1 to trial t is processed as such and can help to detect the asynchrony. This processing of short delays may be impaired in individuals with schizophrenia. It is consistent with previous results, showing higher sensitivity to short asynchronies in individuals with schizophrenia, in spite of the inability to explicitly report them^11,12,34^.

The results are also consistent with our hypothesis that there is a dissociation between the automatic processing of small asynchronies and delays, versus the explicit and conscious processing of time^18^. In individuals with schizophrenia, asynchronies and delays would be detected at the subthreshold level but would not be experienced consciously. Such a difficulty may or may not be related to a general difficulty in conscious realization ^35^. However, whatever the precise mechanisms of the group difference found in the present study, the reduced benefit of SOA increase in individuals with schizophrenia may bear a special clinical significance, if it reveals a lack of explicitly experienced delay from one trial to another. This may correspond to the experience of the individuals with schizophrenia, reporting that time stops and disappears^36^. It may reflect a more general difficulty to experience the passage of time, affecting the processing at the level of seconds as well^20–21^. It may thus contribute to disorders of the sense of self, given the established link between time and self^1–6^.

This study has some limitations. Individuals with schzophrenia were evaluated clinically, but no phenomenological scale was used, limiting our conclusions regarding the clinical meaning of the results. Also our suggestion that the results are related to a general difficulty in the experience of the passage of time is indirect. Future studies should focus on the relationship between the benefit of the passage of time at several time scales, to verify the hypothesis of a lack of interaction between the passage of time at the level of milliseconds and seconds.

## Acknowledgments

The authors thank the University of Strasbourg, the University Hospital of Strasbourg (API-HUS no. 3494), and INSERM, for their support, and the French National Research Agency (Grant ANR-12-SAMA-0016-01) the PHRC (Hospital Project of Clinical Research, PHRCI 2010 − HUS n°4706), and the associations ‘FondaMental’, the ‘Fondation pour la recherche en psychiatrie et en santé mentale’ (http://www.psy-fondation.fr/), as well as ‘APICIL’,for their financial support.

