## Supplemental Information for "Passage of time at the level of milliseconds : a new approach and a selective difficulty in individuals with schizophrenia"

### 1. Regression coefficients for the model of sequential effects on performance

In order to test our hypothesis simultaneously with all three datasets, we used a generalised linear-mixed effect model. Log-odds of correct response were predicted by the fixed factor structure consisting of the group (neurotypical or people with schizophrenia), the study (1-3), the difference in SOA between the two successive trials (decreasing, same or increasing), response on the previous trial (correct or incorrect), and their interactions. A random intercept was specified at the level of participant, nested within the study. However, given the complexity of the model, the full model ran into convergence issues. In order to automate identification of the maximal generalised linear model that converges given the data, we used buildmer[1,2] function, to iterate through the models of different complexity and structure (with fixed and random effects) and to build the model with maximal complexity that is supported by the data (AIC criterion). As shown in the manuscript, the retained model included the four fixed factors, their first order interactions and three three-way interactions. The three-way interactions were found among the difference in SOA, the response on the previous trial and the study; the difference in SOA, the response on the previous trial and group; and the response on the previous trial, group and study. Coefficients of the model are shown in Table 1, and the model structure is indicated above the table. Of note, given the manner in which SOA direction is defined, the three groups were different with respect to SOA on trials t for the three groups. Specifically, in the "increasing" group there were no trials with the smallest, and in the "decreasing" group no trials with the largest SOA (see Fig. S1). Therefore, we excluded from the analyses the two extreme SOAs, in order to match the difficulty across the categories. Results without this exclusion are shown in Fig. S2 and Table S5. Coefficients represent the difference of log-odds between each category and the reference level (conditional on the random effects). For example, the reference level for the variable "group" is individuals with schizophrenia, and the coefficients show the difference between the reference level and the control group (Group = Control). As a measure of effect size, we show odds ratio (OR), indicating a one-unit change in a predictor changes the odds of a correct response on a trial, with values above 1 indicating increased odds and values below 1 indicating decreased odds.[3]

Table 1. Coefficients of the regression model presented in the manuscript.

Response ~ 1 + (1 | Study/Participant) + Study + Previous Response + SOA difference + Study x Previous Response + Group + Previous Response x SOA Difference + SOA difference x Group + Previous Response x Group

| | β (OR) | SE | z | p |
|---|---|---|---|---|
| Intercept | 0.055 (1.056) | 0.147 | 0.376 | 0.707 |
| SOA difference = Increasing | 1.206 (3.340) | 0.058 | 20.635 | 0.000 |
| SOA difference = Same | 0.170 (1.185) | 0.077 | 2.199 | 0.028 |
| Previous response = Correct | 1.403 (4.067) | 0.080 | 17.448 | 0.000 |
| Study 3 | -1.198 (0.301) | 0.183 | -6.553 | 0.000 |
| Study 2 | -0.868 (0.420) | 0.170 | -5.111 | 0.000 |
| Group = Control | 0.266 (1.305) | 0.152 | 1.484 | 0.138 |
| SOA difference = Increasing x Previous response = Correct | -0.185 (0.831) | 0.069 | -2.671 | 0.008 |
| SOA difference = Same x Previous response = Correct | 0.677 (1.968) | 0.086 | 7.862 | 0.000 |
| SOA difference = Increasing x Group = Control | -0.570 (0.565) | 0.070 | 7.411 | 0.000 |
| SOA difference = Same x Group = Control | 0.095 (1.09) | 0.082 | 0.727 | 0.467 |
| Previous response = Correct x Study 3 | 0.487 (1.627) | 0.066 | -8.098 | 0.000 |
| Previous response = Correct x Study 2 | 0.058 (0.059) | 0.080 | 1.158 | 0.247 |
| Previous response = Correct x Group = Control | 0.181 (1.198) | 0.062 | 2.922 | 0.003 |

Table S2. Number of trials per condition in Study 1

| | | **Control participants** | **Individuals with schizophrenia** |
|---|---|---|---|
| | | Median number of trials [Inter-quartile range] | |
| | | With / Without extreme SOA | |
| SOA increasing | $SOA_{t-1}$ incorrect | 59 [30] / 37 [21.5] | 60 [38] / 39 [21] |
| | $SOA_{t-1}$ correct | 67 [34] / 25 [22] | 64 [32] / 23 [23] |
| SOA same | $SOA_{t-1}$ incorrect | 26 [18] / 7 [11] | 31 [27] / 10 [15] |
| | $SOA_{t-1}$ correct | 70 [19] / 41 [9.5] | 61 [23] / 37 [15] |
| SOA decreasing | $SOA_{t-1}$ incorrect | 7 [14] / 5 [5] | 15 [23] / 4 [9] |
| | $SOA_{t-1}$ correct | 114 [17] / 55 [6.25] | 106 [24] / 55 [10] |

Table S3. Number of trials per condition in Study 2

| | | Control participants | Individuals with schizophrenia |
|---|---|---|---|
| | | Median number of trials [Inter-quartile range] | |
| | | With / Without extreme SOA | |
| SOA increasing | $SOA_{t-1}$ incorrect | 43 [15] / [30 [11] | 55.5 [15] / 35.5 [9.75] |
| | $SOA_{t-1}$ correct | 38 [12] / 19 [8] | 26.5 [8.75] /12 [7.5] |
| SOA same | $SOA_{t-1}$ incorrect | 21 [12] / 9.5 [5.25] | 30 [11.5] / 15 [7.5] |
| | $SOA_{t-1}$ correct | 55 [12] / 36 [6] | 45 [13.8] / 28 [12] |
| SOA decreasing | $SOA_{t-1}$ incorrect | 6 [6] / 2[1] | 14.5 [10] / 5 [6] |
| | $SOA_{t-1}$ correct | 74 [6] / 47 [3] | 66 [7.75] / 44 [6.75] |

Table S4. Number of trials per condition in Study 3

| | | Control participants | Individuals with schizophrenia |
|---|---|---|---|
| | | Median number of trials [Inter-quartile range] | |
| | | With / Without extreme SOA | |
| SOA increasing | $SOA_{t-1}$ incorrect | 181 [60.5] / 157 [52.5] | 195 [71] /166 [55] |
| | $SOA_{t-1}$ correct | 118 [67.2] / 87.5 [53] | 90 [77] / 69 [61.5] |
| SOA same | $SOA_{t-1}$ incorrect | 26[13.8] / 21 [12.75] | 29 [22] / 21 [18] |
| | $SOA_{t-1}$ correct | 26 [8.5] / 21 [10.25] | 24 [17] / 20 [13.5] |
| SOA decreasing | $SOA_{t-1}$ incorrect | 79 [60.5] / 56.5 [52.5] | 91 [62] / 63 [53] |
| | $SOA_{t-1}$ correct | 228 [67.8] / 194.5 [49.25] | 200 [98] / 175 [84] |

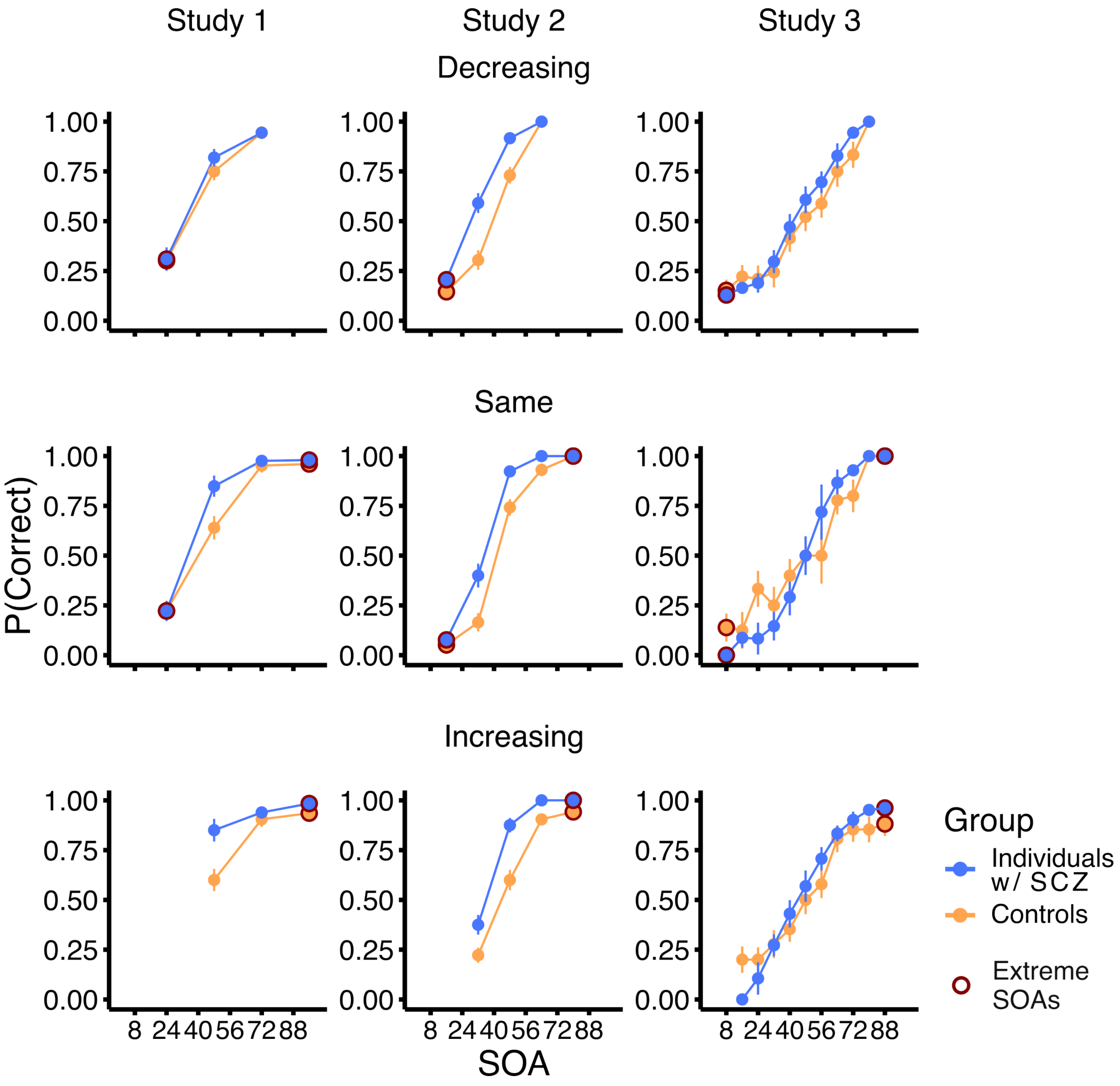


Figure S1. Average performance in the three studies. The average performance is plotted as a function of SOA on that trial, for the three studies (columns) and the two groups (color coded). Performance is shown separately for the three categories of the difference between the SOA on trial t (on the x-axis) and the previous SOA (t-1). The SOA either decreased (top), was the same (middle) or increased (bottom) from one trial to the next. Red symbols indicate the extreme SOAs that are not shared between the three groups, and were excluded from the analysis presented in the manuscript.

**2. Re-analysis with inclusion of the extreme SOAs**

When all trials were included in the analysis (even those with the extreme SOAs), there was a significant effect of the SOA direction ($\chi^2(2) = 1766.640$, $p<0.01$), and the response on the previous trial ($\chi^2(1) = 377.240$, $p<0.01$). Relative to the condition where the SOA decreased from trial t-1 to trial t the performance was better when the SOA increased ($\beta_{\text{SOA Increasing}} = 1.904$, SE = 0.05, $z = 33.745$, $p < 0.01$,

OR = 6.712) and when it remained the same ($\beta_{SOA\ Same}$ = 0.052, SE = 0.071, z = 0.738, p = 0.461, OR = 1.053). Furthermore, when the response on the previous trial was correct, log-odds of correct response on trial t were greater ($\beta_{Response\_t\text{-}1=Correct}$ = 1.076, SE = 0.055, z = 19.423, $p < 0.01$, OR = 2.933).

There was no significant main effect of the group ($\chi^2(1)$ = 0.025, p=0.873). Importantly, the group interacted with the SOA direction ($\chi^2(2)$ = 83.315, p<0.01), such that performance of participants in the control group increased more when the SOA increased from the previous to the current trial, relative to individuals with schizophrenia ($\beta_{SOA\ Increasing\ x\ Control}$ = 0.568, SE = 0.088, z = 6.445, $p < 0.01$, OR = 1.765; $\beta_{SOA\ Same\ x\ Control}$ = -0.108, SE = 0.110, z = -0.982, p = 0.326, OR = 0.898).

The SOA direction also interacted with the previous response ($\chi^2(2)$ = 350.410, p<0.01). The interaction with the previous response indicated that an increase in performance for trials with a relative when SOA from one trial to the next repeated, the increase in the performance greater when response on the previous trial was correct ($\beta_{SOA\ Same\ x\ Response\_t\text{-}1=Correct}$ = 1.384, SE = 0.087, z = 15.833, $p < 0.01$, OR = 3.991; $\beta_{SOA\ Increasing\ x\ Response\_t\text{-}1=Correct}$ = -0.1322, SE = 0.076, z = -1.742, p = 0.081, OR = 0.876). There was also evidence for an interaction between the previous response and the group ($\chi^2(1)$ = 4.1, p<0.01), suggesting that the effect of the previous response was stronger in the control group ($\beta_{Response\_t\text{-}1=Correct\ x\ Group\ =\ Control}$ = 0.173, SE = 0.085, z = 2.025, $p < 0.05$, OR = 1.189).

We also found evidence for a three-way interaction. There was evidence for an interaction between the SOA direction, response on the previous trial and the group ($\chi^2(2)$ = 7.586, p<0.01). These interactions reflect the intricacies of the effects among the three studies, and further details are shown in Table S5.

In summary, as in the previous analysis, we found a complex pattern of effects between the tested variables. Importantly, even when all trials were included, there was an interaction between the group and the SOA direction, indicating the robustness of the results. The same pattern of results was also found when each experiment was examined separately.

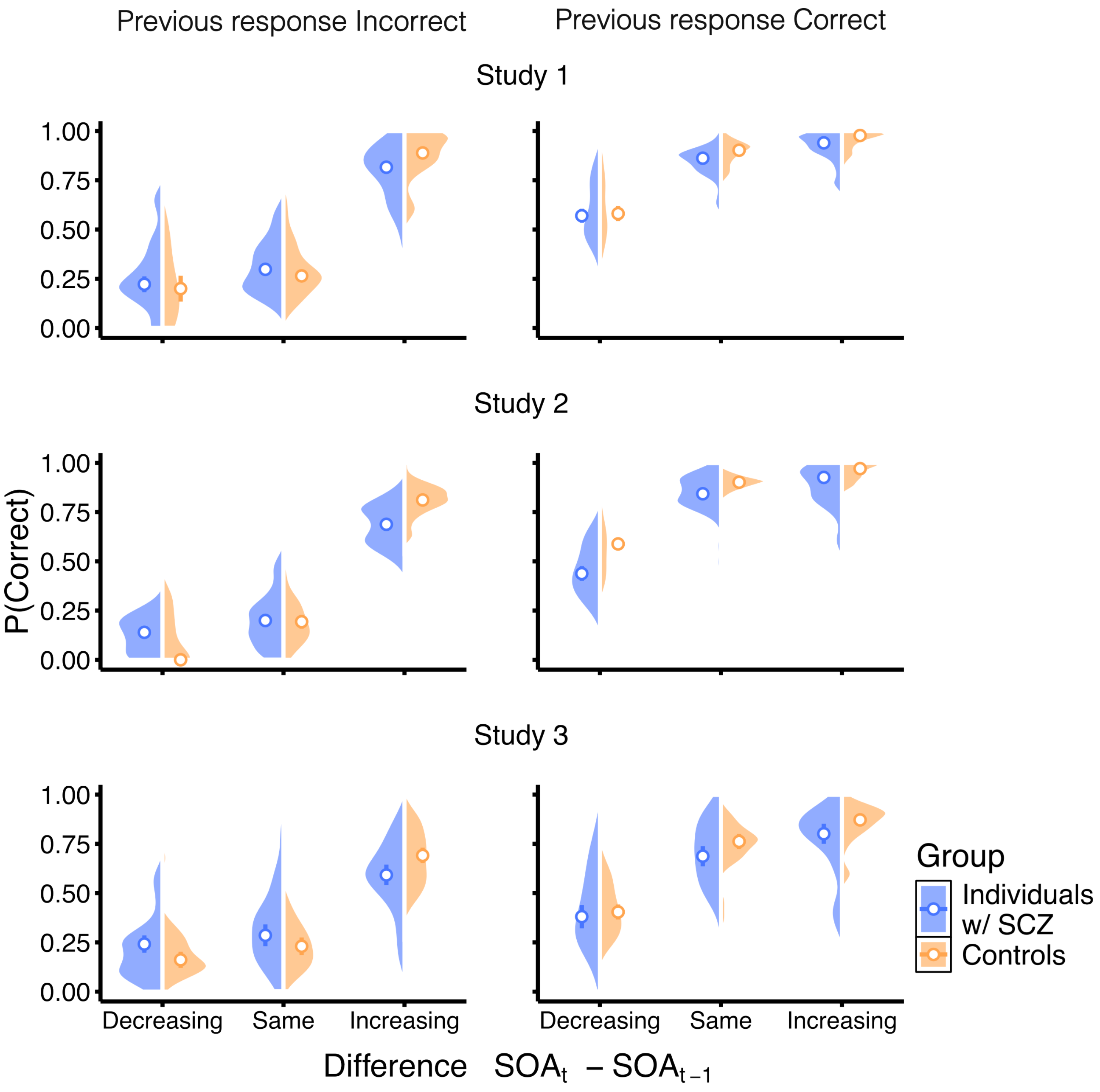


Figure S2. Sequential effects in the asynchrony detection task. Proportion of correct responses is plotted as a function of the difference between the SOA on the previous (t-1) and current (t) trial. The differences between the two consecutive trials are grouped in three groups, with respect to direction of SOA change (Decreasing, Same or Increasing). The left and right columns show performance when response on the previous trial (t-1) was incorrect and correct, respectively. The rows show performance for the three analysed datasets. Color codes the group, with blue symbols indicating performance of individuals with schizophrenia and orange that of individuals in the control group. Violin plots show distributions of the proportions of correct responses, for the two groups, for all trials (including the two extreme SOAs). Circles show median performance (error bars indicate standard error of the median).

Table S5. Coefficients of the regression model fit to the entire dataset.

Response ~ 1 + (1|Study/Participant) + SOA difference + Previous Response + Group + SOA difference x Previous Response + SOA difference x Group + Previous Response x Group + SOA difference x Previous Response x Group

| | β (OR) | SE | z | p |
|---|---|---|---|---|
| Intercept | -1.023 (0.359) | 0.094 | -10.893 | 0.000 |
| SOA difference = Increasing | 1.905 ( 6.719) | 0.056 | 33.745 | 0.000 |
| SOA difference = Same | 0.052 (1.053) | 0.071 | 0.738 | 0.461 |
| Previous response = Correct | 1.076 (2.933) | 0.055 | 19.423 | 0.000 |
| Group = Control | 0.022 (1.022) | 0.137 | 0.160 | 0.873 |
| SOA difference = Increasing x Previous response = Correct | -0.132 (0.876) | 0.076 | -1.742 | 0.082 |
| SOA difference = Same x Previous response = Correct | 1.386 (3.999) | 0.088 | 15.833 | 0.000 |
| SOA difference = Increasing x Group = Control | 0.568 (1.765) | 0.088 | 6.445 | 0.000 |
| SOA difference = Same x Group = Control | -0.108 (0.898) | 0.110 | -0.982 | 0.326 |
| Previous response = Correct x Group = Control | 0.173 (1.189) | 0.085 | 2.025 | 0.043 |
| SOA difference = Increasing x Previous response = Correct x Group = Control | 0.002 (1.002) | 0.116 | 0.021 | 0.983 |
| SOA difference = Same x Previous response = Correct x Group = Control | 0.323 (1.381) | 0.132 | 2.449 | 0.014 |

#### 3. Sequential effects with inclusion of the SOA on the current trial

In analyses shown above, the performance was grouped across different levels of SOA on the current trial (t). This decision was made in order to simplify the analyses and maximize the number of trials for each condition. Nevertheless, we have also examined the observed sequential effects with SOA on the current trial. Figures S3 and S4 visualise performance as a function of SOA on the current trial and Table S6 shows the results of a regression model that best describes the data.

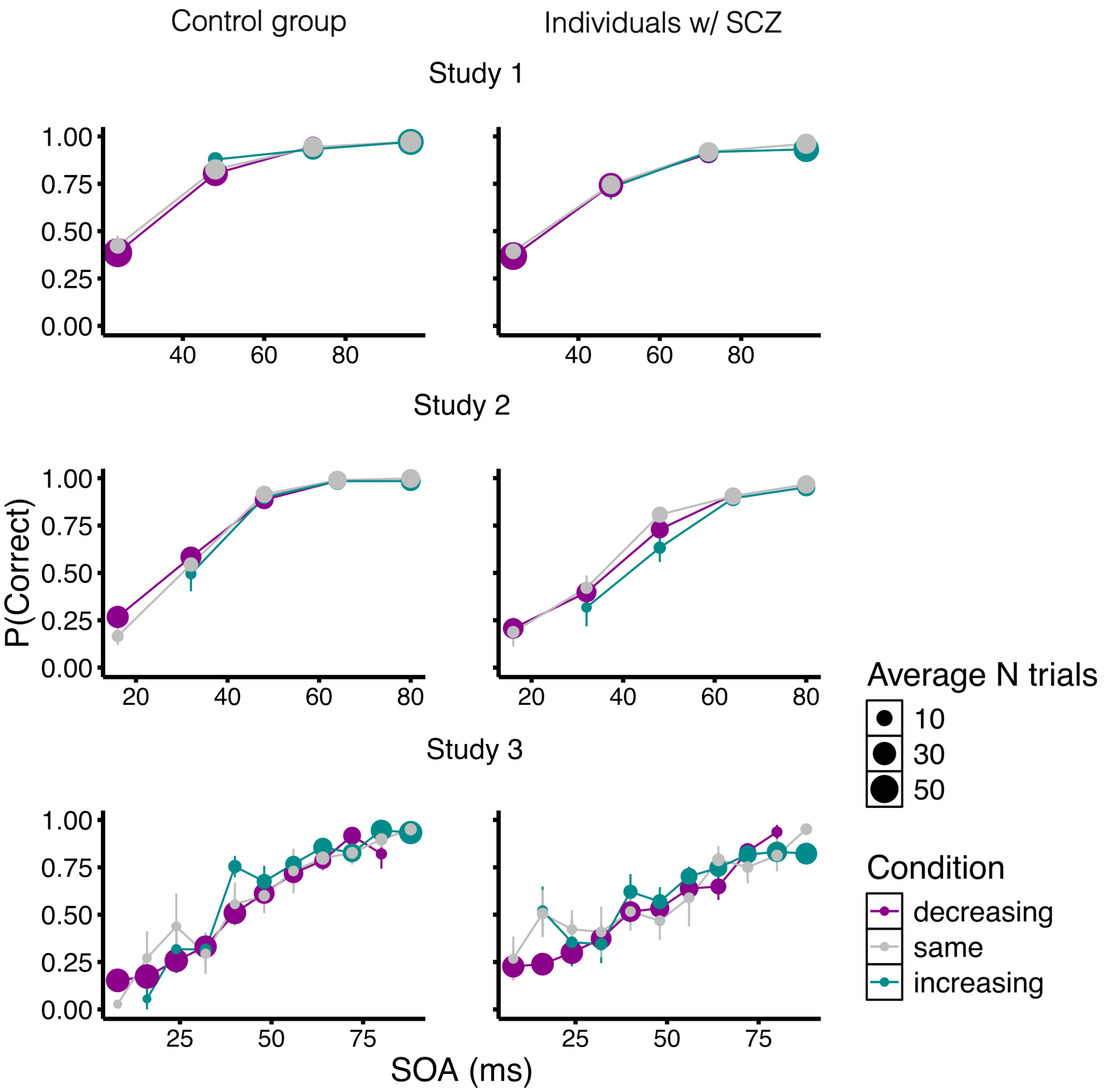


Figure S3. Sequential effects in the asynchrony detection task, when response on previous trial was correct. Proportion of correct responses is plotted as a function of the SOA on the current (t). The differences between the two consecutive trials are grouped in three groups, with respect to direction of SOA change (Decreasing, Same or Increasing) and color coded. The left and right columns show performance for the two groups. The rows show performance for the three analysed Studies. Symbols show average performance and error bars indicate standard errors of the mean between participants. Size of the symbols corresponds to the average number of trials for each condition.

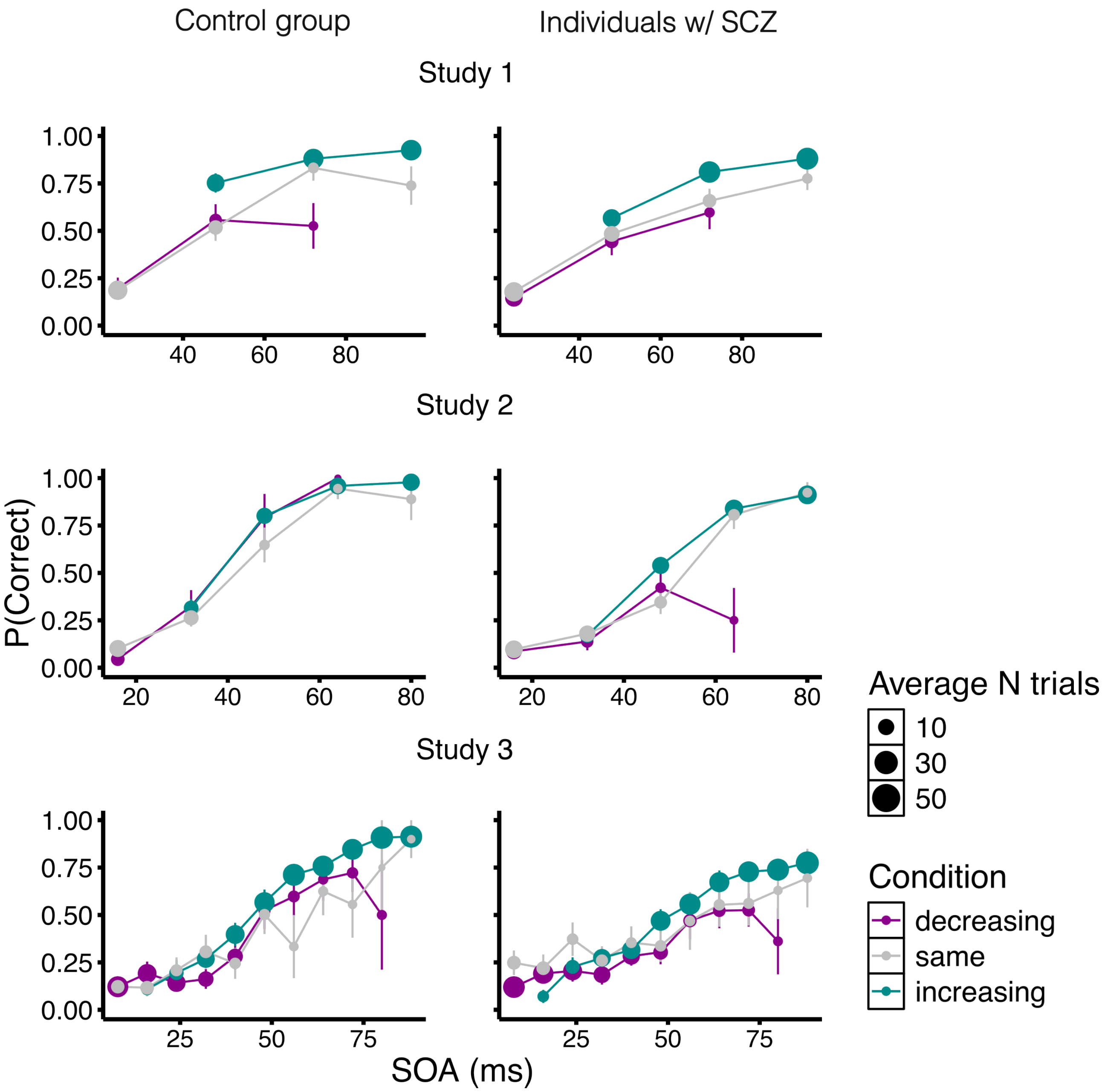


Figure S4. Sequential effects in the asynchrony detection task, when response on previous trial was incorrect. As in Figure S3, proportion of correct responses is plotted as a function of the SOA on the current (t). The differences between the two consecutive trials are grouped in three groups, with respect to direction of SOA change (Decreasing, Same or Increasing) and color coded. The left and right columns show performance for the two groups. The rows show performance for the three analysed Studies. Symbols show average performance and error bars indicate standard errors of the mean between participants. Size of the symbols corresponds to the average number of trials for each condition.

Table S6. Coefficients of the regression model that included SOA on the current trial

Response ~ 1 + (1|Study/Participant) + SOA difference + Previous Response + Study + Group + SOA Difference x Previous Response + Previous Response x Study + SOA Difference x Group + Previous Response x Group

| | β (OR) | SE | z | p |
|---|---|---|---|---|
| Intercept | 0.055 (1.056) | 0.147 | 0.376 | 0.707 |
| SOA difference = Increasing | 1.206 (3.340) | 0.058 | 20.635 | 0.000 |
| SOA difference = Same | 0.170 (1.185) | 0.077 | 2.199 | 0.028 |
| Previous response = Correct | 1.403 (4.067) | 0.080 | 17.448 | 0.000 |
| Study 3 | -1.198 (0.301) | 0.183 | -6.553 | 0.000 |
| Study 2 | -0.868 (0.420) | 0.170 | -5.111 | 0.000 |
| Group = Controls | 0.226 (1.254) | 0.152 | 1.484 | 0.138 |
| SOA difference = Increasing x Previous response = Correct | -0.185 (0.831) | 0.069 | -2.671 | 0.008 |
| SOA difference = Same x Previous response = Correct | 0.677 (1.968) | 0.086 | 7.862 | 0.000 |
| Previous response = Correct x Study 3 | -0.570 ( 0.565) | 0.070 | -8.098 | 0.000 |
| Previous response = Correct x Study 2 | 0.095 (1.1) | 0.082 | 1.158 | 0.247 |
| SOA difference = Increasing x Group = Controls | 0.487 (1.627) | 0.066 | 7.411 | 0.000 |
| SOA difference = Same x Group = Controls | 0.058 (1.06) | 0.080 | 0.727 | 0.467 |
| Previous response = Correct x Group = Controls | 0.181 (1.198) | 0.062 | 2.922 | 0.003 |

**4. Study 4 – Masking experiment - Different types of masking**

In the masking experiment (Study 4) we did not find evidence for an interaction between the group and the direction of SOA change from the previous to the current trial on the performance, but we did find evidence for a main effect of the SOA direction change. We also analysed separately the condition where the type of masking was the same from one trial to the next. Results are shown in Table S7 and S8.

Table S7. Coefficients of the regression model in which only trial that had identical masking (backward-backward or forward-forward) were considered
Response ~ 1 + (1|Participant) + SOA difference + Previous Response + Group + Previous Response x Group

| | β (OR) | SE | z | p |
|---|---|---|---|---|
| Intercept | -0.156 (0.856) | 0.172 | -0.905 | 0.366 |
| Previous response = Correct | -0.083 (0.920) | 0.129 | -0.643 | 0.520 |
| SOA difference = Increasing | 0.284 (1.328) | 0.096 | 2.956 | 0.003 |
| SOA difference = Same | 0.089 (1.093) | 0.133 | 0.670 | 0.503 |
| Group = Controls | -0.092 (0.912) | 0.234 | -0.394 | 0.693 |
| Previous response = Correct x Group = Controls | 0.825 (2.282) | 0.182 | 4.533 | 0.000 |

Table S8. Coefficients of the regression model in which only trial that had different masking (backward-forward or backward-forward) were considered
Response ~ 1 + (1|Participant)

| | β (OR) | SE | z | p |
|---|---|---|---|---|
| Intercept | 0.035 (1.036) | 0.115 | 0.307 | 0.759 |

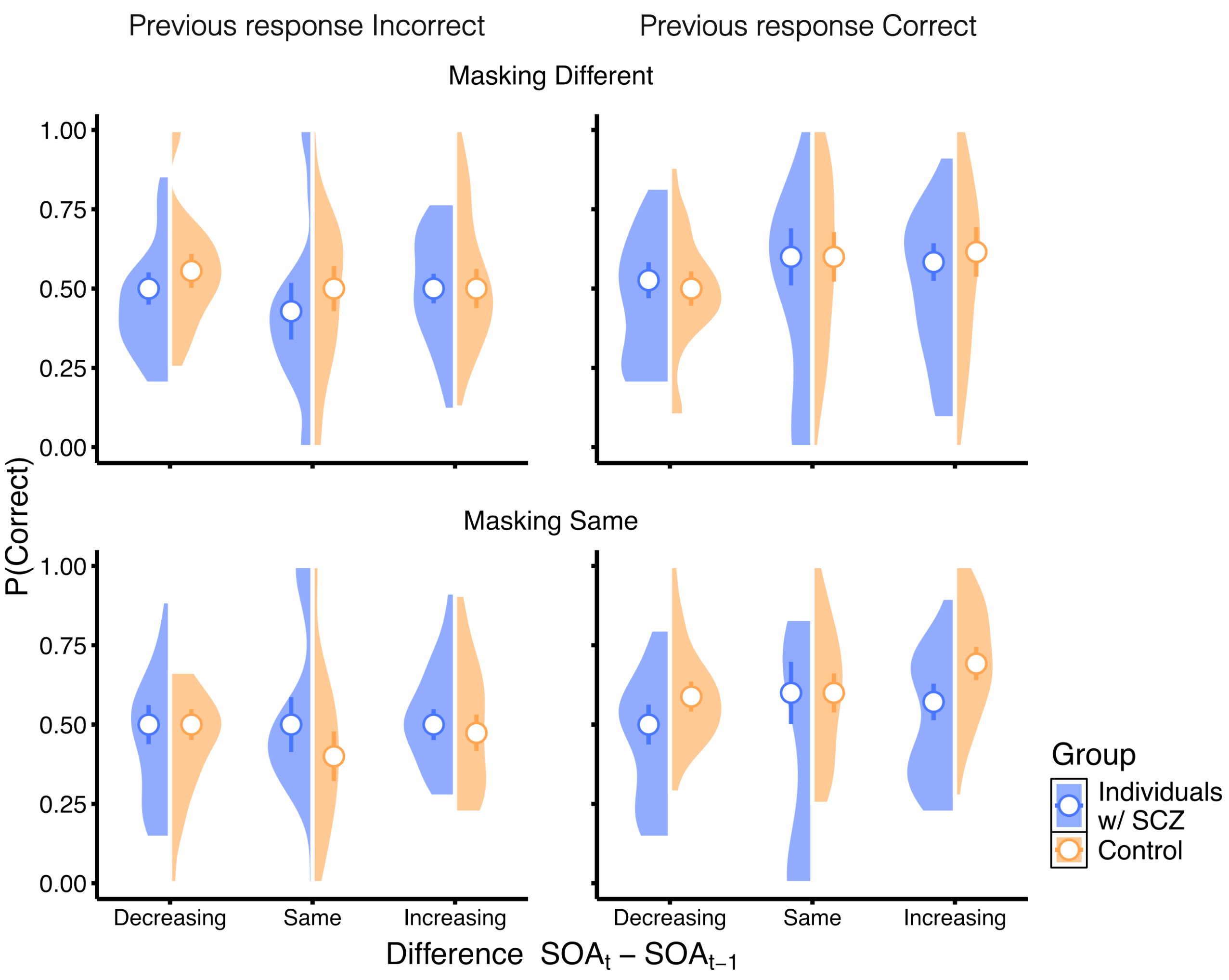


Figure S5. Sequential effects in the masking task. Proportion of correct responses is plotted as a function of the difference between the SOA on the previous (t-1) and current (t) trial. The differences between the two consecutive trials are grouped in three groups, with respect to direction of SOA change (Decreasing, Same or Increasing). The left and right columns show performance when response on the previous trial (t-1) was incorrect and correct, respectively. The rows show performance for the relative masking types between trials t-1 and t (Same if masking on both trial t-1 and t were either backward or forward, and different otherwise). Color codes the group, with blue symbols indicating performance of individuals with schizophrenia and orange that of individuals in the control group. Violin plots show distributions of the proportions of correct responses, for the two groups. Circles show median performance and error bars indicate standard error of the median.

**5. Masking with inclusion of extreme SOAs**

Results of the masking data presented in the main manuscript were done only on intermediate SOAs (12, 18 and 24 ms), in order to match the difficulty of the task for the three categories of the relative SOA difference (SOA decreasing, staying the same or increasing). We have also done the same analysis with inclusion of all trials (except when SOA was 0). The results were comparable, indicating that the effects are robust to small differences in difficulties on trial t. Regression coefficients of the retained model are shown in Table S9.

Table S9. Coefficients of the regression model for the masking experiment – extreme SOAs included

Response ~ 1 + (1|Participant) + SOA difference + Previous Response + Group + SOA difference x Previous Response + Previous Response x Group + Masking Type + Previous Response x Masking Type + Gropu x Masking Type + Previous Response x Group x Masking Type

| | β (OR) | SE | z | p |
|---|---|---|---|---|
| Intercept | -0.384 (0.681) | 0.152 | -2.530 | 0.011 |
| SOA difference = Increasing | 0.349 (1.418) | 0.072 | 4.816 | 0.000 |
| SOA difference = Sames | 0.008 (1.001) | 0.097 | 0.084 | 0.933 |
| Previous Response = Correct | -0.023 (0.977) | 0.109 | -0.207 | 0.836 |
| Group = Control | 0.361 (1.435) | 0.209 | 1.728 | 0.084 |
| Masking Type = Same | 0.056 (1.058) | 0.090 | 0.621 | 0.535 |
| SOA difference = Increasing x Previous Response = Correct | 0.256 (1.292) | 0.103 | 2.481 | 0.013 |
| SOA difference = Same x Previous Response = Correct | 0.225 (1.252) | 0.136 | 1.652 | 0.098 |
| Previous Response = Correct x Group = Control | -0.152 (0.859) | 0.134 | -1.139 | 0.255 |
| Previous Response = Correct x Masking Type = Same | -0.115 (0.891) | 0.132 | -0.875 | 0.382 |
| Group = Control x Masking Type = Same | -0.178 (0.837) | 0.132 | -1.346 | 0.178 |
| Previous Response = Correct x Group = Control x Masking type = Same | 0.548 (1.730) | 0.187 | 2.929 | 0.003 |

### 6. Masking - Analyses by the group

In order to verify that both groups did benefit from the (e.g. that the effect of the SOA difference is not driven by an improvement in the control group), we repeated the analysis separately for the two groups. Results are shown in Tables S10 and 11.

Table S10. Coefficients of the regression model for the masking experiment – control group only

Response ~ 1 + (1|Participant) + Previous Response + SOA difference + Masking Type + Previous Response x SOA difference + Previous Response x Masking Type

| | β (OR) | SE | z | p |
|---|---|---|---|---|
| Intercept | 0.219 (1.244) | 0.186 | 1.179 | 0.238 |
| Previous response = Correct | -0.214 (0.807) | 0.158 | -1.356 | 0.175 |
| SOA difference = Increasing | -0.024 (0.976) | 0.138 | -0.175 | 0.861 |
| SOA difference = Same | -0.225 (0.798) | 0.182 | -1.239 | 0.215 |
| Masking type = Same | -0.237 (0.789) | 0.125 | -1.889 | 0.059 |
| Previous response = Correct x SOA difference = Increasing | 0.366 (1.442) | 0.195 | 1.877 | 0.061 |
| Previous response = Correct x SOA difference = Same | 0.510 (1.665) | 0.246 | 2.073 | 0.038 |
| Previous response = Correct x Masking type = Same | 0.654 (1.923) | 0.174 | 3.755 | 0.000 |

Table S11. Coefficients of the regression model for the masking experiment – individuals with schizophrenia only

Response ~ 1 + (1|Participant) + SOA difference

| | β (OR) | SE | z | p |
|---|---|---|---|---|
| Intercept | -0.164 (0.848) | 0.61 | -1.018 | 0.309 |
| SOA difference = Increasing | 0.212 (1.236) | 0.093 | 2.293 | 0.022 |
| SOA difference = Same | 0.079 (1.082) | 0.125 | 0.604 | 0.546 |
